# Cryo-electron microscopy structure of a zinc uptake ABC transporter

**DOI:** 10.1101/2025.07.03.663024

**Authors:** Changxu Pang, Hoang Nguyen, Qingfang Zhang, Ivet Bahar, Qun Liu

## Abstract

Zn^2+^ is an essential micronutrient to all living organisms and plays a key role in various physiological functions. Microorganisms employ high-affinity Zn^2+^ ABC transporters to uptake zinc from the environment when it is scarce. However, the mechanism of zinc uptake and its regulation remain poorly understood. Here, we report the cryo-electron microscopy structure of the Zn^2+^ ABC transporter complex ZnuB-ZnuC from *Escherichia Coli*. The complex has two ZnuB subunits for transport and two ZnuC subunits for regulation. The ZnuB homodimer is in an outward-facing, closed conformation with a large hydrophilic cavity at the dimer interface. The ZnuC subunits contain an N-terminal ATP-binding cassette (ABC) domain (NBD) and a C-terminal zinc-sensing domain (ZSD). Zn^2+^ binding to the ZSD locks the transporter in a closed state regardless of nucleotide, whereas under low-Zn^2+^ conditions, ZSD C-terminal disorder permits ATP-driven zinc uptake. High-affinity zinc ABC transporters are ubiquitously utilized by pathogenic bacteria to evade host immune systems in competing for essential Zn^2+^. These findings highlight new pharmaceutical targets for disrupting Zn^2+^ homeostasis in antibiotic-resistant pathogens.

**One-sentence summary:** High-affinity zinc ABC transporter structures show a self-regulating mechanism via a built-in zinc sensor.

## Introduction

The zinc ion (Zn^2+^) is an essential micronutrient supporting all living organisms. It plays physiological functions in gene transcription, structure rigidity, development, photosynthesis, immune response, and numerous Zn^2+^-dependent enzymatic activities ^1^. Zn^2+^ is impermeable to membranes and has to be transported across cell membranes by Zn^2+^ uptake transporters, including Zrt-/Irt-like proteins (ZIPs) ^2,3^, ATP-binding cassette (ABC) transporters^4^, and natural resistance-associated macrophage proteins (NRAMP)^5^ before being used by cells. While Zn^2+^ is an essential micronutrient, nevertheless, its excessive accumulation is cytotoxic, and its intracellular concentration has to be tightly regulated. As revealed by Zn^2+^ uptake assays, Zn^2+^ ABC transporters are high-affinity transporters whose expression levels are tightly regulated by the Zn^2+^-sensing transcription factor Zur ^6^. Many bacterial pathogens use ABC transporters to compete for Zn^2+^ with their hosts - humans, animals, and plants. Conversely, upon sensing a pathogen attack, the hosts produce Zn^2+^-chelating proteins and molecules to limit the access of Zn^2+^ to be used by pathogens, forming an adaptive immunity called nutritional immunity ^7-16^. Antibiotic resistance is a growing global health crisis. A recent analysis reported that AMR caused 1.27 million deaths directly in 2019, in addition to contributing an additional 5 million indirect deaths^17^. These pathogen-specific ABC transporters may be attractive targets for the development of novel antibiotics and vaccines. However, the structure and mechanism of function of Zn^2+^ ABC transporters, and how their activities are regulated and linked to intracellular Zn^2+^ contents, remain largely unclear.

ABC transporters are a large family of transporters ubiquitous among all living organisms. They use ATP hydrolysis to drive the transport of ions, metabolites, proteins, nucleotides, and other biological molecules across membranes ^18,19^. Many human ABC transporters are also important drug targets for diseases ^18^. The ABC transporters contain at least two ATP-binding cassette (ABC) domains (also called nucleotide-binding domains, NBDs) and two transmembrane (TM) domains (TMD). The NBD regulates the transport activity in the TMD through ATP binding and hydrolysis. ZnuABC is an *E. coli* ABC transporter; it is composed of five subunits: two transmembrane (ZnuB) subunits, two ATP-binding cassette (ABC) regulatory (ZnuC) subunits, and one Zn^2+^ binding protein substrate (ZnuA)^20^.

In this work, we present the cryo-electron microscopy (cryoEM) structures of a high-affinity Zn^2+^ ABC transporter ZnuB-ZnuC, later referred to as ZnuBC tetrameric complex from *E. coli*. The structures are resolved in outward-facing Zn^2+^-bound as well as Zn^2+^-free states. We identify a Zn^2+^ sensing domain (ZSD) at the C-terminus of the regulatory subunit ZnuC. ZSD detects intracellular Zn^2+^ content and regulates the transport activity. Our structural, functional, and computational analyses of the tetrameric ZnuBC reveal a mechanism of Zn^2+^ uptake and regulation through a built-in intracellular Zn^2+^ sensor.

## Results

### Structure determination

The expression of *ZnuA, ZnuB, and ZnuC* in *E. coli* is regulated by the *zur* transcription factor as illustrated in **Figure S1A**. We cloned the *E. coli ZnuA and ZnuBC* genes to the pETDuet1 vector with one cassette containing *ZnuBC* and the other cassette containing *ZnuA* (**Figure S1B**). We expressed the complex in bacteria, purified it in detergent, and reconstituted it into amphipol nanodiscs using a workflow summarized in **Figure S1C**. The complex contained ZnuB and ZnuC (ZnuBC), lacking ZnuA (**Figure S1D-E**). We determined its structure using single-particle cryo-electron microscopy (CryoEM). To obtain the structure, we prepared the ZnuBC particles (**Figure S2A**) in the presence of adenylyl-imidodiphosphate (AMP-PNP), a non-hydrolyzing ATP analog. Averaged 2D classes showed multiple views of the ZnuBC complex (**Figure S2B**). The reconstructed 3D map shows a belt of amphipol densities around the transporter (**Figure S2C**). The cryoEM data analysis workflow is summarized in **Figure S3**. The reconstructed cryoEM map used 34,187 particles, reaching a resolution of 2.5 Å (**Figure S4A**). The final reconstruction contains particles with a widespread distribution in Euler space with slightly more views at elevation angles around zero (**Figure S5A**). The cryoEM density is of high quality, showing all nine transmembrane helices and most side-chain features in a transporter subunit of ZnuB (**Figure S6**).

### Overall structure

The ZnuBC structure is a heterotetramer, consisting of two copies of ZnuB and ZnuC subunits (**Figure 1A-B**). ZnuB is the zinc transporter subunit. The two ZnuB subunits form a dimer with an interface of 1503 Å^2^ as calculated by PISA ^21^. Each ZnuB monomer contains nine TM helices (TM1-TM9) and two short α-helices: α1 in the periplasm and α2 in the cytosol (**Figure 1B-C**). Most TM helices are bent or tilted inside the membrane. TM5 is long and extends from the membrane into the cytosol, followed by α2, which is known as a coupling helix in ABC transporters ^19^. TM6 and TM7 do not fully cross the membrane; they form a kink inside the membrane (**Figure 1C**). The intramembrane loop between TM6 and TM7, L(6,7), has close contact with the periplasmic helix α1, which connects to TM4 and TM5 through two long loops (**Figure 1C**).

**Figure 1.**
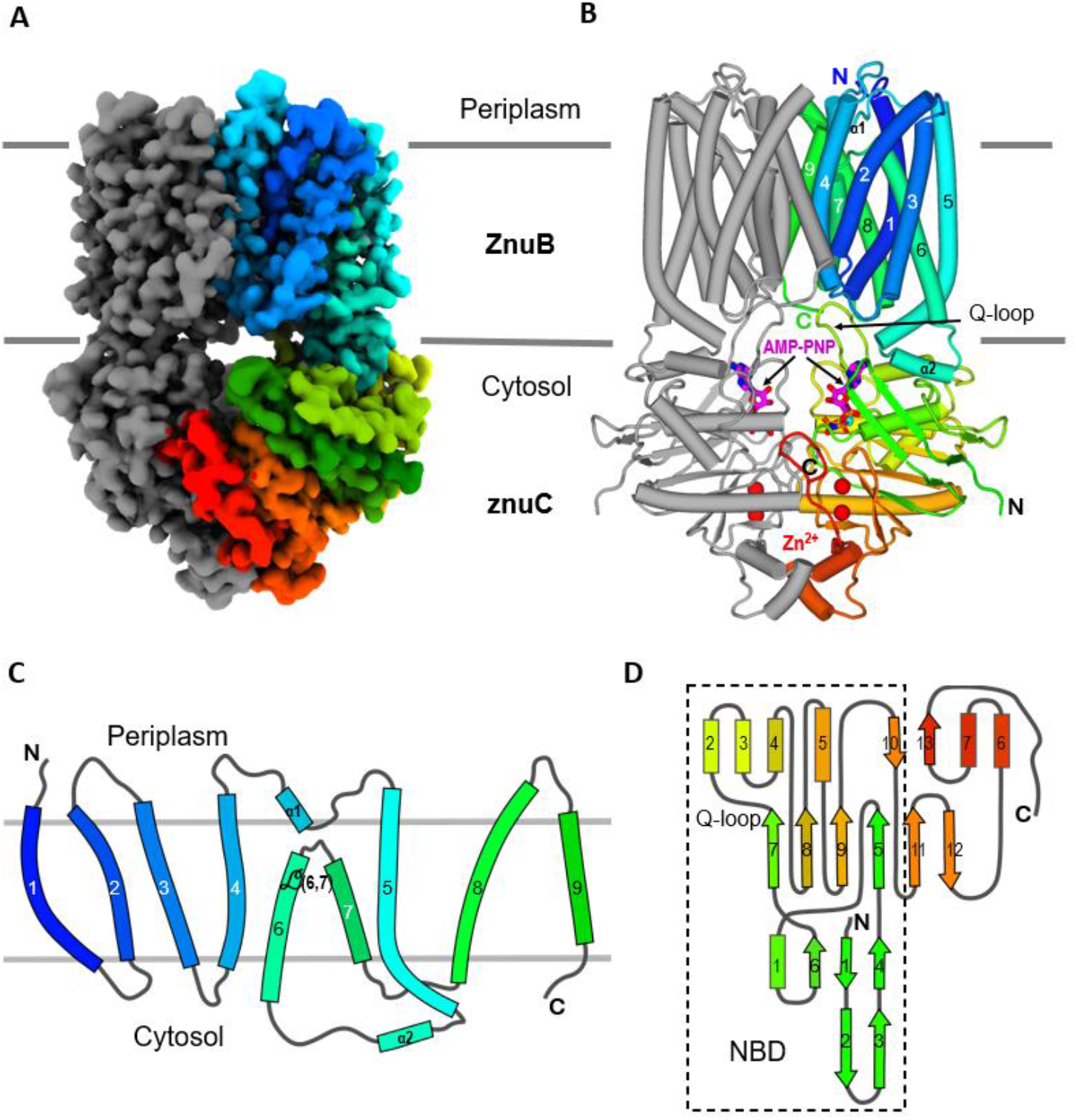
Overall structure of the ZnuB-ZnuC (ZnuBC) transporter. (**A)** CryoEM density map of the ZnuBC subunits of the ZnuABC transporter. One subunit is colored rainbow, and the other, gray. **(B)** Overall structure of the ZnuBC dimer. The coloring is as in panel a. Four-bound Zn^2+^ ions are shown as red spheres. Two AMP-PNP molecules are shown as magenta sticks. **(C)** Topology diagram of a ZnuB subunit. (**D)** Topology diagram of a ZnuC subunit.

ZnuC is the regulatory subunit, located in the cytosol. It interacts with ZnuB mainly through the α2 on ZnuB and a Q-shaped loop (Q-loop) on ZnuC (**Figure 1B**). The two ZnuC subunits form a dimer with an interface of 1606 Å^2^. Each ZnuC monomer contains an AMP-PNP molecule bound to its NBD domain. In addition to the NBD domain, the C-terminus of ZnuC has an extended domain consisting of three β-strands (β11-13), two short α-helices (α6 and α7), and a long loop (**Figure 1D**). Surprisingly, the extension harbors two Zn^2+^ ions in each of the ZnuC subunits (**Figure 1B**).

### ZnuB zinc transporter

The ZnuB dimer has a two-fold symmetry; TM2, TM4, TM7, and TM9 from each monomer form the dimer interface (**Figure 2A-B**). A 40 Å-long transport channel from the periplasmic side to the cytosol is at the dimer interface (**Figure 2C-D**), lined by TM2 and TM4 residues of each subunit. These two helices on each ZnuB subunit bend to expose a vestibule to the periplasm with the α1 helices inserted as a plug to constrict the channel. Therefore, we interpret this structure as an outward-facing, closed state. The channel is wide in the middle and has two restriction bottlenecks. On the periplasmic side, the bottleneck is formed by the hydrophobic residues Met112, Leu115, and Phe116 from α1 and Leu190 from the loop L(6,7). On the cytosol side, the bottleneck is formed mainly by Tyr33 residues from both subunits.

**Figure 2.**
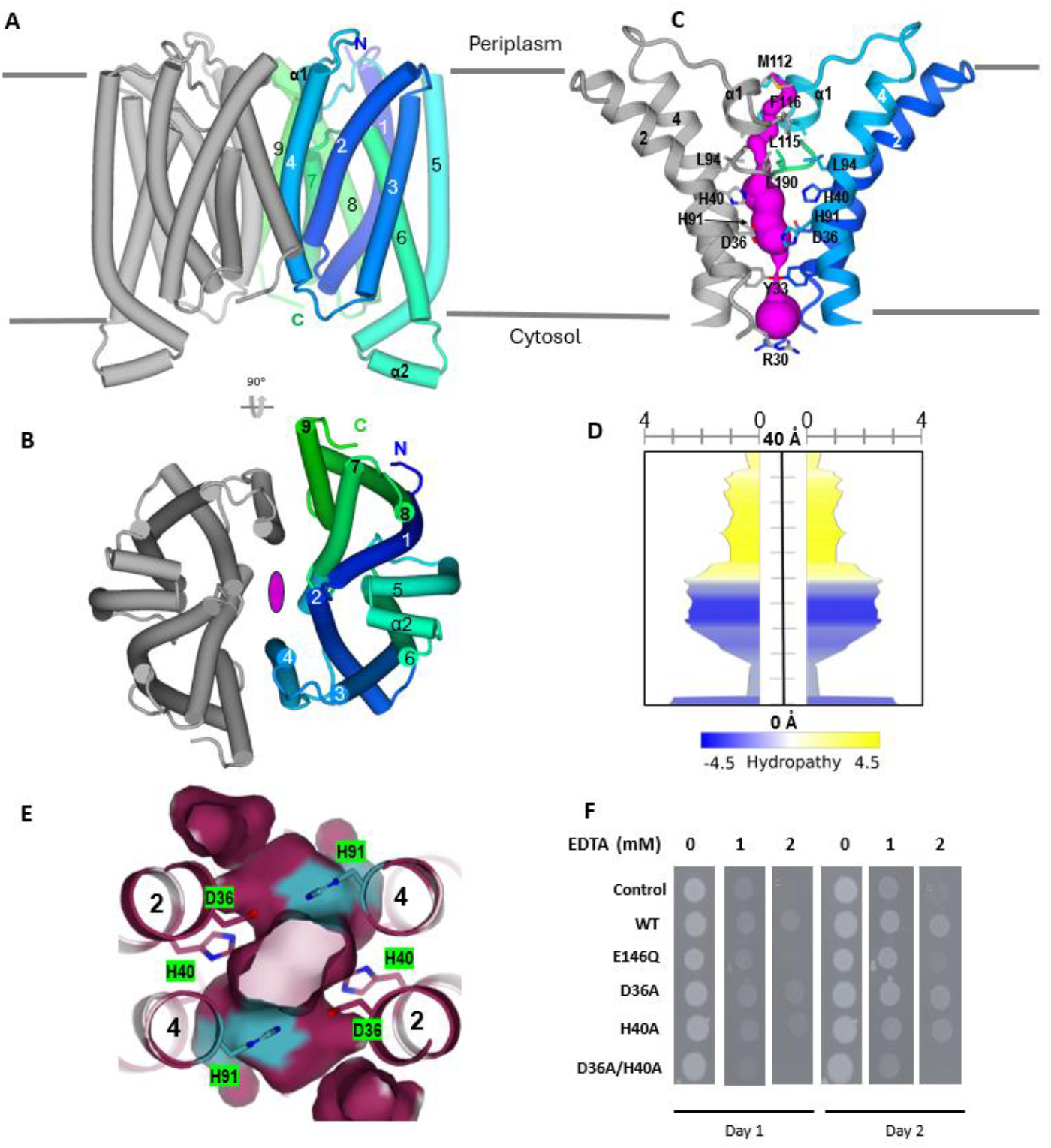
Structure of ZnuB dimer. **(A-B)** Two views of the overall structure of the dimer, viewed from the side and the cytosol, respectively. The dimer has a twofold symmetry. (**C**) The membrane transport channel *(magenta surface)* lined by the labeled TM2 and TM4 residues from each monomer. (**D**) Quantification of the size and hydrophobicity of the transport channel. (**E**) An elongated cavity at the center of the transmembrane region formed by histidine and aspartate residues. The surface color shows the conservation level: *magenta*, more conserved; *cyan*, relatively less conserved. (**F**) Functional characterization of zinc uptake and cell viability by removing Zn^2+^ using EDTA.

Between the two bottlenecks, there is a large cavity of about 13 Å long and 6 Å wide. The cavity is hydrophobic on the periplasmic side, formed by residues Leu94 and Leu190 (**Figure 2C-D**). The hydrophobic residues may help seal the transporter to prevent a leaky uptake of Zn^2+^. The cytoplasmic side is hydrophilic, formed by conserved residues Asp36, His40, and less conserved residue His91 (**Figures 2E and S7**). The cavity is elongated and has enough space to accommodate one Zn^2+^ inside. We hypothesized that the side chains of the conserved Asp36 and His40 might contribute to the Zn^2+^ transport activity. We thus produced D36A, H40A, and D36A-H40A mutants, transformed them with bacterial cells, and measured cell growth viability under Ethylenediaminetetraacetic acid (EDTA) treatment (**Figures 2F and S8A**). EDTA treatment removed Zn^2+^ from uptake by bacteria. Compared to the plasmid control and wild-type (WT), the D36A-H40A double mutant did not grow well under Zn^2+^-limiting conditions, treated with 2 mM EDTA, suggesting their essential role in Zn^2+^ uptake. To test if His91 is involved in the ZnuB transport function, we produced H91A, D36A-H91A, and H40A-H91A mutants and performed cell viability assays in the presence of various concentrations of EDTA (**Figure S8B**), we found that cells overexpressing H91A grown similarly to the wild-type; D36A-H91A and H40A-H191A double mutants remain active in Zn^2+^ uptake, suggesting that His91 is not as important as Asp36 and His40 in Zn^2+^ transport. In addition, bacterial cells transformed with WT grew better than the control under the Zn^2+^-limiting condition, implicating that ZnuB was expressed and contributed to the uptake of Zn^2+^ from the Zn^2+^-limiting environment.

### ZnuC nucleotide-binding domain

ZnuC is the regulatory subunit of the Zn^2+^ ABC transport system. The solved AMP-PNP-bound ZnuC dimer contains two AMP-PNP molecules and four Zn^2+^ ions (**Figure 3A-B**). The AMP-PNP molecule binds at the N-terminus of α1, which is sandwiched by two β-sheets. In other ABC transporters, ATP or AMP-PNP binding triggers dimerization through an ATP ^22^. However, in the ZnuC dimer structure, AMP-PNP has no interactions with the other ZnuC subunit, indicating not involved in dimerization (**Figures 3A and S9A)**.

**Figure 3.**
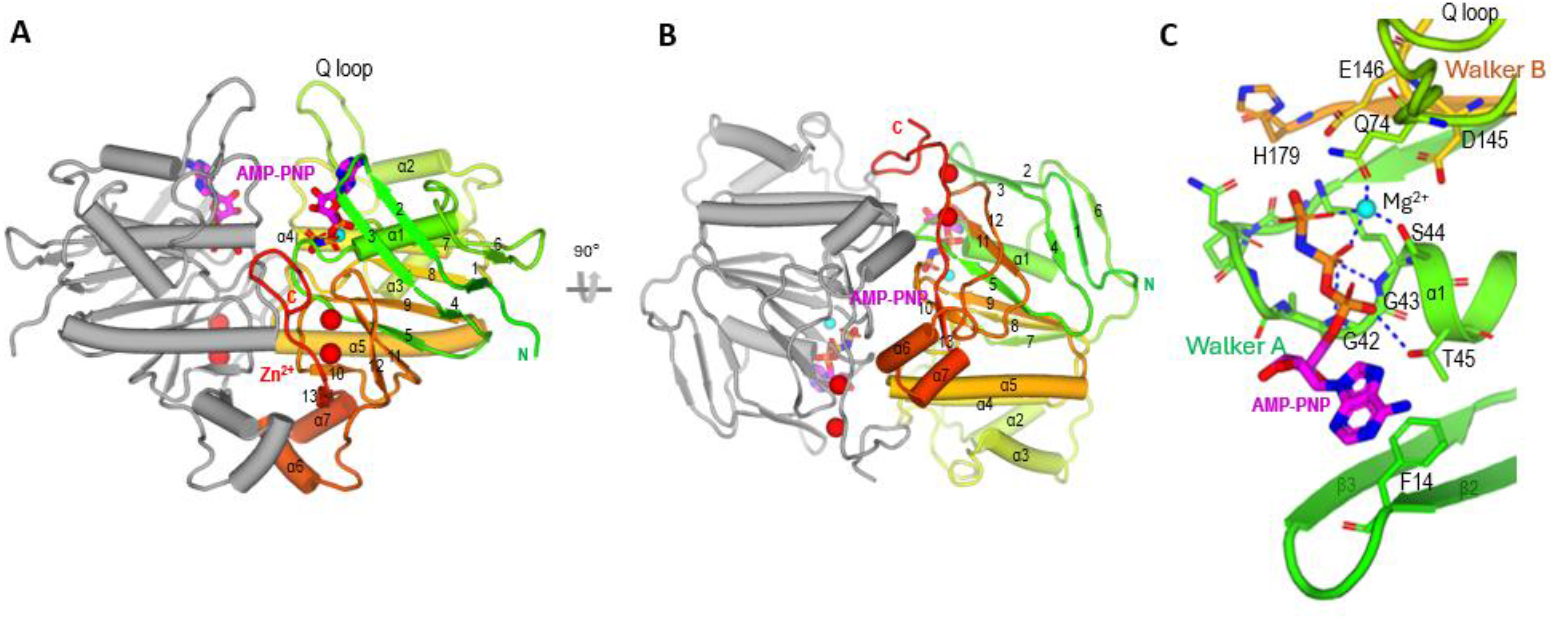
Structure of the ATP-binding cassette subunit ZnuC. **(A-B)** Two views of the AMP-PNP-bound structure of the ZnuC dimer. The coloring is the same as **Figure 1B**. (**C**) Interactions between AMP-PNP and the NBD of ZnuC. The *cyan sphere* is an Mg^2+^ ion, coordinated by S44, Q74, and AMP-PNP.

AMP-PNP binds to the NBD primarily via the Walker A motif (**Figure 3C**). Its phosphate groups form hydrogen bonds with Gly42, Gly43, and Thr45, while Ser44 and two phosphate oxygens coordinate the Mg^2+^ ion that is also liganded by Glu74 in the Q-loop (residues 74–85). The side chain of the Walker B catalytic residue, Glu146, points directly at the γ-phosphate, positioning it for hydrolysis. Meanwhile, the adenine ring of AMP-PNP π-stacks against Phe14 in β2.

We also produced an ATPase-deficient ZnuBC E146Q mutant and solved its cryoEM structure in the presence of ATP at 2.9 Å resolution (**Figures S4B and S5B**). The solved structure is very similar to the AMP-PNP-bound structure; neither ATP is involved in the dimerization of ZnuC (**Figure S9B**). All of these contacts shown in the AMP-PNP binding are retained in the ATP-bound E146Q mutant structure (**Figure S9B**).

To assess nucleotide-binding induced conformational changes, we solved an apo-form, nucleotide-free ZnuBC structure (**Figures S4C and S5C**) and overlaid it with the AMP-PNP–bound structure. ZnuB, including the coupling helix α2 (residues 154-162), shows virtually no shift (RMSD 0.337 Å over 260 ZnuB Cα atoms), and ZnuC also remains largely unchanged (RMSD 0.455 Å over 242 ZnuC Cα atoms). The notable conformational changes localize to the nucleotide pocket (**Figure S9C**). In the ZnuC apo structure, the sidechain of Gln74 in the Q-loop reoriented due to the loss of interaction with the Mg^2+^. However, the Q-loop backbone itself is unaltered. Phe14 in β2 and the associated β-sheet shifted due to the loss of the *π* stacking with the nucleotide adenine ring. In the ZnuB apo structure, there is a movement of the L(3,4) loop (residues 73-82), likely due to its interactions with the β-sheet in ZnuC. Compared with the apo structure, ATP or AMP-PNP binding at the Walker A site does not trigger significant global rearrangements in either ZnuB or ZnuC.

### ZnuC Zn^2+^ binding and autoregulation

In both the apo and nucleotide-bound ZnuBC structures, the C-terminus of each ZnuC monomer harbors two Zn^2+^ sites, M1 and M2 (**Figure 4A**). M1 is coordinated by Cys193, Cys200, His230 and His232; M2 by His197, Cys199, His234 and His246. These two Zn^2+^ ions lock the ZnuC C-terminus into an ordered conformation, and helices α6–α7 from each monomer pack tightly to form a dimer interface burying ∼807 Å^2^ (**Figure 3A**). This Zn^2+^-stabilized dimer interface via helices α6–α7 might prevent the ATP-induced global conformational switch, confining movement to local changes around the nucleotide pocket (**Figure S9C**).

**Figure 4.**
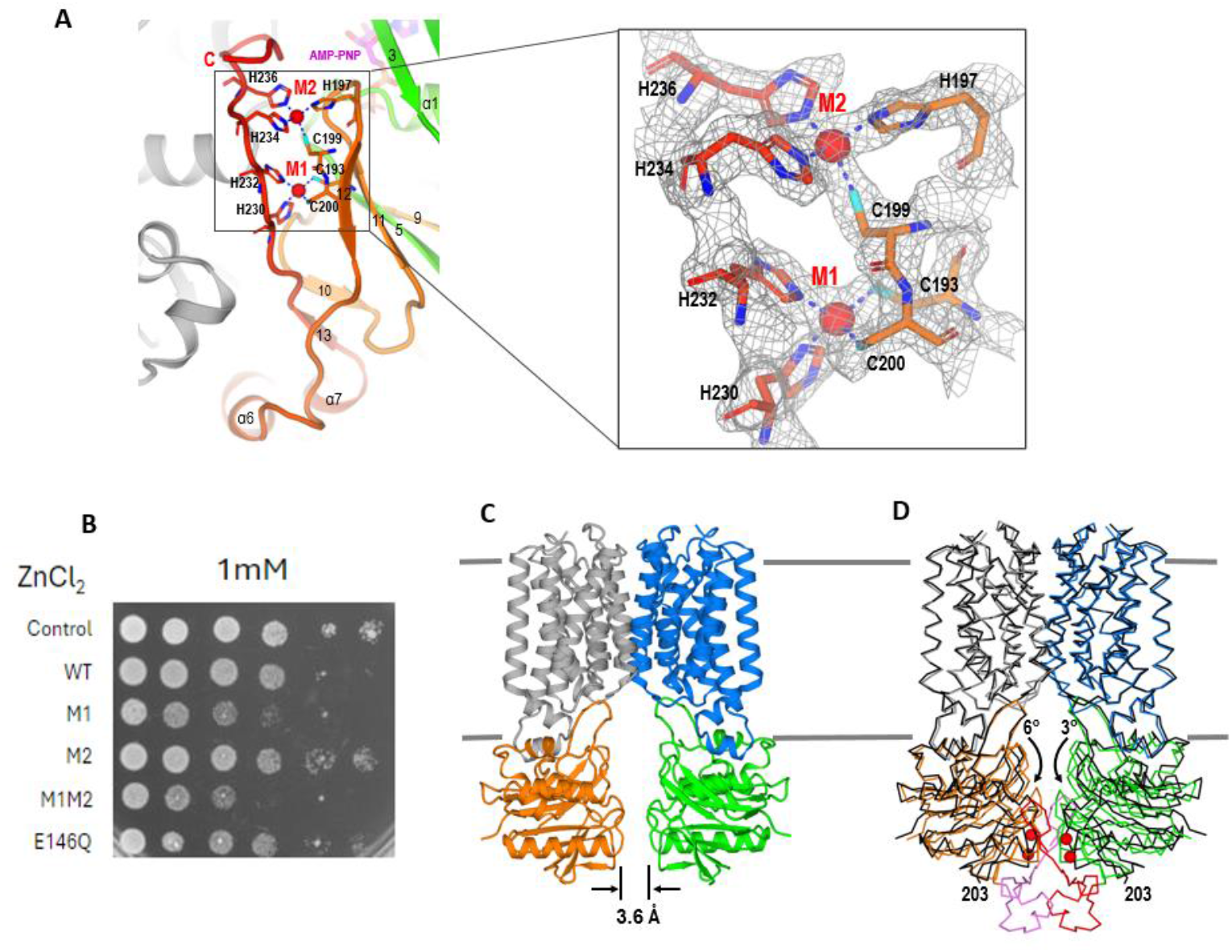
Intracellular zinc-sensing and transport cycle regulation by ZnuC ZSD. (**A**), Zn^2+^ binding sites M1 and M2 at the ZnuC ZSD. Inet shows the cryoEM density for the M1 and M2 sites. (**B**) Cell growth viability assays of ZnuBC wild-type and mutants. *E. coli* C43 cells transformed with the indicated plasmids for plating Argar-LB plates with buffer or 1 mM Zn^2+^ and the presence of 0.2 mM IPTG (final). Compared with the control, which has an empty plasmid, M1 or M1M2 double mutants in ZnuC showed cell toxicity. (**C**) CryoEM structure of EDTA-treated Zn^2+^-free state. Compared to the apo structure, the two ZnuC monomers moved away from each other by 3.6 Å. (**D**) Superimposed apo and ETDA-treated structures to show conformational changes associated with Zn^2+^ sensing and regulation. Arrows show rotations of ZnuC relative to the respective interacting ZnuB.

We therefore asked whether M1 and M2 sites regulate Zn^2+^ uptake. We replaced each coordinating Cys/His with alanine, expressed the mutants in *E. coli*, and compared growth in media containing Zn^2+^ (**Figures 4B and S10**). The cells expressing M2 mutant behaved like wild type, but the cells expressing M1 or M1M2 double mutants showed markedly reduced viability under 1 mM Zn^2+^, consistent with the loss of regulation, leading to intracellular Zn^2+^ overloading and toxicity. We therefore hypothesize that the C-terminal domain of ZnuC (residues 193–251) functions as an intracellular Zn^2+^ sensor and designate it the zinc-sensing domain (ZSD).

Based on the structure of the M1 and M2 sites, we identified a strictly conserved motif, Cys193xxxHisxxCysCys204, in 5,712 ZnuC homologs, including major human and plant pathogens, using PatternSearch^23^ (**Figure S11**). Alignment of eight representative pathogens confirmed absolute conservation of all M1/M2 residues (**Figure S12**), suggesting that Zn^2+^-mediated ZnuC dimerization is a common mechanism for controlling zinc uptake and homeostasis across these pathogenic bacteria.

Because Zn^2+^-binding stabilizes the ZnuC zinc-sensing domain (ZSD) structure, we hypothesized that Zn^2+^ removal would destabilize it. To test this, we treated ZnuBC with EDTA to strip Zn^2+^ and solved its cryoEM structure (**Figures S4D and S5D**). The EDTA-treated structure lacked density for most of the ZSD and lost all Zn^2+^ ions (**Figure 4D-E**). Alignment with the apo structure revealed broken two-fold symmetry: ZnuC subunits rotated by 6° and 3° relative to their ZnuB partners, reducing the ZnuC–ZnuC interface from 1,606 Å^2^ to 86 Å^2^, while ZnuB remained largely unchanged (RMSD 0.660 Å) (**Figure 4E**).

These structural and functional data suggest that Zn^2+^ depletion unfolds ZSD and increases ZnuC flexibility by disrupting its dimer interface. We hypothesize that ZSD may sense and monitor intracellular Zn^2+^ content and autoregulate the transporter function to prevent over-accumulation of cellular Zn^2+^.

### Alternating access transport model

To explore Zn^2+^ transport mechanisms in ZnuBC, we employed anisotropic network model (ANM) analysis ^24,25^ on cryoEM structures. While molecular dynamics simulations offer atomic detail, they are limited by sampling and timescale, especially for multimeric proteins in membranes. ANM-based normal mode analysis, via the ProDy interface ^26^, efficiently reveals global, cooperative motions. We computed intrinsic dynamics for apo and EDTA-treated ZnuBC, both isolated and in explicit lipid bilayers, using the membrANM extension^27^. The four lowest-frequency modes (ANM1–4) reflect the transporter’s architecture and are largely unaffected by the membrane (**Figure S13A-B**). ANM1 involves global twisting of ZnuB and ZnuC dimers; ANM2, bending of ZnuB relative to ZnuC; ANM3, tilting; and ANM4, alternating periplasmic/cytosolic opening (**Figure S13C**). Mode spectra comparison shows ANM1, ANM2, and ANM4 are conserved (cosine similarity >0.80) despite Zn^2+^-induced changes. ANM4 is particularly insightful, predicting large-scale conformational shifts, enabling Zn^2+^ access from the periplasm and release into the cytosol, consistent with an alternating-access model ^28^. (**Figure S13D**). Transitions involve exposure and narrowing of the hydrophobic cavity (TM2/TM4) and coordinated rotation of the α2 helix and ZnuC dimer rearrangement, facilitating Zn^2+^ release. However, ANM does not resolve localized motions, which likely occur at higher frequencies.

## Discussion

Excess Zn^2+^ is toxic, so its uptake and intracellular levels must be tightly controlled. In this work, we determined cryoEM structures of a high-affinity Zn^2+^ uptake ABC transporter ZnuBC in its nucleotide/Zn^2+^, apo/Zn^2+,^ and apo/Zn^2+^-free states. The identification of a Zn^2+^ sensing domain (ZSD) at the C-terminus of ZnuC, together with the functional characterization of the M1 and M2 sites, allowed us to propose an autoregulation mechanism for Zn^2+^ uptake and homeostasis. Under Zn^2+^-replete conditions, each ZnuC zinc-sensing domain (ZSD) binds Zn^2+^, dimerizes, and locks ZnuB in an outward-facing, closed conformation that is refractory to ATP or AMP-PNP binding (**Figure 3A**). When cellular Zn^2+^ is depleted by enzymes or transcription factors, Zn^2+^ dissociates from the ZSDs, destabilizing their dimer interface and disordering the ZnuC NBDs (**Figure 4C**), allowing the transporter to use ATP to drive the uptake of Zn^2+^ from periplasm.

ZnuABC expression is controlled by the Zur transcription factor, but its post-translational regulation was unknown. We discovered a Zn^2+^ sensor in the C-terminal zinc-sensing domain (ZSD) of ZnuC, defined by the conserved motif CxxxHxCC. This motif creates two high-affinity Zn^2+^ sites (M1 and M2) that prevent Zn^2+^ overaccumulation and support cell growth (**Figure 4A-B**). Based on the classification of ABC transporters ^29^, ZnuABC belongs to type II ABC transporters. Five other type II ABC transporter structures have been reported: ButCD-F ^30^ (PDB code 2QI9), HI1470/1^31^ (PDB code 2NQ2), FhuCDB ^32^ (PDB code 7LB8), BhuUV ^33^ (PDB code 5B57), and PsaBC ^34^ (PDB code 7KYP). We performed structural alignments of the ZnuC ZSD with the NBD C-termini of the five ABC importers (PsaC, ButD, HI1470, FhuD, and BhuV), as shown in **Figure S14**. Unlike the ZnuC ZSD, which comprises two short α-helices (α6-7), a β-strand (β13), and a long loop, the C-termini of the compared NBDs consist of either a single long helix (PsaC, **Figure S14B**) or a long helix followed by two β-strands (ButD, HI1470, FhuD, and BhuV, **Figure S14C-F**). These differences highlight the unique structural features of the ZnuC ZSD in Zn^2+^ sensing and uptake regulation.

Beyond the Zn^2+^ ABC transporters, we previously proposed that bacterial ZIP-family Zn^2+^ transporter *Bb*ZIP has a histidine-rich loop between TM3 and TM4 and functions as an intracellular Zn^2+^ sensor to regulate the transporter activity ^35^. In plants, a histidine-rich loop in Arabidopsis IRT1, also a ZIP member, senses metals and triggers endosomal sorting and vacuolar degradation of IRT1 ^36^. A histidine-rich cluster in human ZIP4 mediates its ubiquitination and degradation to prevent Zn^2+^ toxicity ^37^. It appears that all living organisms have evolved versatile ways to regulate the activity of Zn^2+^ transporters post-translationally through inhibition or degradation to prevent excess accumulation of Zn^2+^ and toxicity. Further studies are needed to see if this regulation exists in other essential but toxic metal micronutrients.

Zn^2+^ ABC transporters are bacterium-specific and exist in most high-risk pathogens (**Figures S11 and S12**). The structures of the Zn^2+^ ABC transporter ZnuBC provide a basis for the development of antibiotics and vaccines targeting the conserved sites, such as the transmembrane cavity in the ZnuB dimer interface. The cavity is elongated and larger than accommodating a single Zn^2+^. A potential inhibitor blocking the cavity may help reduce pathogenicity by preventing the evasion of pathogens from hosts’ nutritional immunity.

## Lead contact

Requests for further information should be directed to the lead contact, Qun Liu.

## Materials availability

All materials produced in this study will be made available upon reasonable request.

## Data and materials availability

The 3-dimensional cryo-EM density maps have been deposited in the Electron Microscopy Data Bank under the accession numbers EMD-70570 for WT-AMP-PNP/Zn^2+^ state [https://www.emdataresource.org/EMD-70570], EMD-70571 for E146Q-ATP/Zn^2+^ state [https://www.emdataresource.org/EMD-70571], EMD-70572 for apo/Zn^2+^ state [https://www.emdataresource.org/EMD-70572], EMD-70573 for apo/Zn^2+^-free state [https://www.emdataresource.org/EMD-70573]. Atomic coordinates have been deposited in the Protein Data Bank under the accession numbers 9OKO for WT-AMP-PNP/Zn^2+^ state [https://doi.org/10.2210/pdb9oko/pdb], 9OKP for E146Q-ATP/Zn^2+^ state [https://doi.org/10.2210/pdb9okp/pdb], 9OKQ for apo/Zn^2+^ state [https://doi.org/10.2210/pdb9okq/pdb], and 9OKR for EDTA-treated, Zn^2+^-free state [https://doi.org/10.2210/pdb9okr/pdb].

## Acknowledgments

We thank the LBMS staff for their help with the cryo-EM operation and data acquisition. This work was supported by the U.S. Department of Energy (DOE), Office of Biological and Environmental Research KP1601011. Partial support from NIH R01GM139297 is gratefully acknowledged by I.B. The work used the Laboratory for Biomolecular Structure (LBMS), which is supported by the U.S. Department of Energy, Office of Science, Office of Biological and Environmental Research.

## Author contributions

Q. Liu designed the study and experiments. C. Pang, Q. Zhang, H. N., and Q. Liu performed the experiments. C. Pang, I. Bahar, and Q. Liu analyzed the data. Q. Liu and I. Bahar wrote the manuscript with help from other coauthors.

## Methods

### Materials and Methods ZnuA-ZnuB-ZnuC cloning

The *E*.*coli* gene *ZnuCB* was amplified from genomic DNA (strain DH5α) using the forward primer 5′-GCCAGGATCCGAATTCACAAGTCTGGTTTCCCTGGA-3′ and the reverse primer 5′-GGCGAAGCTTTTAGCTGGCCTGCTTTTTCA-3′, which introduced unique BamHI and HindIII restriction sites flanking the *ZnuCB* gene. The *E. coli ZnuA* was amplified from genomic DNA (strain DH5α) using the forward primer 5′-TATACATATGTTACATAAAAAAACGCTTCT -3′ and the reverse primer 5′-CAGACTCGAGATCTCCTTTCAGGCAGCTCG -3′, which introduced unique NdeI and XhoI restriction sites flanking the ZnuA gene. The PCR products were inserted into the pGEM-T-Easy vector according to the manufacturer’s instructions. The sequences were confirmed by DNA sequencing with the sequencing primers M13F 5′-GTAAAACGACGGCCAGTG-3’ and M13R 5’-GGAAACAGCTATGACCATG-3’. The *ZnuCB* was digested with BamHI and HindIII and ligated into the pET-Duet1 vector. Then *ZnuA* was digested with NdeI and XhoI and ligated into the pET-Duet1-ZnuCB. The sequence was confirmed by DNA sequencing with sequencing. All primers used are listed in **Table S2**. The assembled plasmid pET-Duet1_ZnuA-ZnuCB was then transformed into *E. coli* C43 (DE3) cells.

### ZnuB-ZnuC expression and purification

The cells transformed with pET-Duet1_ZnuA-ZnuCB were cultured at 37℃ to the density an OD600 of 0.8-1.2 in Terrific Broth (TB) medium supplemented with 100 μg/ml carbenicillin. Protein production was induced with 0.5 mM isopropyl-β-D-thiogalactopyranoside (IPTG). Cells were harvested 4 hrs after the IPTG induction by centrifugation at 6000xg for 15 min. Cells were resuspended in cold lysate buffer (20 mM HEPES, pH 7.4, 200 mM NaCl, and 20% glycerol) containing 0.1 mM phenylmethysulfonyl fluoride (PMSF). The cells were lysed by passage through an EmlsiFlex-C3 homogenizer (Avestin, Ottawa, Canada) six times at 15,000 psi. Cell debris was removed by centrifugation 31,360xg for 1 hr. The supernatant was collected, and the membrane was harvested by centrifugation at 223,608xg for 2 hrs. The membrane pellet was resuspended in resuspend buffer (20 mM HEPES, pH 7.4, 200 mM NaCl, 20% glycerol, and 10 mM imidazole) supplemented with protease inhibitor cocktail EDTA-free (abcam, Waltham, MA, Cat# ab270055). The membrane was solubilized with 1.5% n-Dodecyl-B-D-Maltoside (DDM) supplemented with 50 μM ZnCl2 for 2 hrs. The insoluble components were removed by centrifugation at 113,210xg for 1 hr. The supernatant was applied to a pre-equilibrated Ni^2+^-NTA column (Anatrace, Maumee, OH, Cat# SUPER-NINTA100). The column was washed by 10 column volumes (CV) of wash buffer (20 mM HEPES, pH 7.4, 200 mM NaCl, 20% glycerol, 40 mM imidazole, and 0.05% DDM). The protein was eluted by elute buffer (20 mM HEPES, pH 7.4, 200 mM NaCl, 200 mM imidazole, and 0.05% DDM) and concentrated to 5 mg/ml by using a 100 kDa molecular-weight cutoff concentrator (Millipore Sigma, Burlington, MA, USA). The purified complex contained ZnuBC (**Fig. S1**).

### Amphipol reconstitution of the ZnuB-ZnuC complex

The concentrated protein (5mg/ml) was mixed with amphipol A8-35 (Anatrace, Maumee, OH, Cat# A8-35) in a mass ratio of protein/amphipol of 1:5 (w/w). The mixture was incubated at 4℃ for 20 hrs. Then, SM-2 Bio-Beads (Bio-Rad, Cat#1523920) were added at a mass ratio of protein/beads of 1:100 (w/w) to the mixture. The mixture was incubated at 4℃ for 3 hrs. Prior to use, Bio-Beads were soaked with buffer containing 20 mM HEPES, pH 7.4, and 200 mM NaCl overnight at 4℃. The Bio-Beads were removed by passing through a micro Bio-Spin chromatography column. The flow-through containing the reconstitution mixture was centrifuged at 20,000xg for 1 hr at 4℃. The supernatant was applied to a Superdex 200 Increase column (GE Healthcare, Inc., Chicago, IL, USA) and eluted in a buffer containing 20 HEPES, pH 7.4, and 100 mM NaCl. The fractions containing the ZnuBC complex were collected and concentrated to 2 mg/ml for cryo-electron microscopy analysis. To prepare the EDTA-treated ZnuBC sample, the amphiphilic-reconstituted ZnuBC sample was incubated with 0.5 mM EDTA for 2 hr on ice before loading onto the pre-equilibrated Superdex 200 column.

### Bacterial growth viability assay

The mutation expression constructs were created with Q5 Site-Directed Mutagenesis Kit (New England BioLabs, Inc., Ipswich, MA) according to the manufacturer’s instructions. The primers used are listed in **Table S1**. The empty pETDuet-1 vector was used for vector control.

The expression constructs were transformed into *E. coli* C43 (DE3) competent cells. Three colonies were inoculated into 5 mL of LB media at 37°C for overnight culture. 100 μl culture was used to inoculate 5 ml of fresh LB media. When the OD600 value reached between 0.8-1.0, 1 mM IPTG (final) was added to induce the expression at 37 °C for 2 hours. Cells were then diluted to OD600 of 0.1, 0.01, 0.001, 0.0001, 0.00001, and 0.000001 with dilution buffer (20 mM HEPES, pH 7.4, 135 mM NaCl, and 5 mM KCl). Two microliters of diluted cells were placed on LB-Agar plates supplemented with 100 μg/ml carbenicillin, 1 mM IPTG, and various concentrations of ethylenediaminetetraacetic acid (EDTA, 0, 1, 2, and 3 mM), ZnCl2 (0, 1, 2 mM), and their combinations. The plates were placed at 37 °C for bacterial growth. The plates were imaged every 24 hrs over three days.

### CryoEM sample preparation and data collection

Three microliters of the reconstituted nanodiscs containing nucleotide-free ZnuBC were applied to a glow-discharged (15 mA current for 15 sec) 300-mesh R 0.6/1 UltrAuFoil Holey Gold grid (Electron Microscopy Sciences, Hatfield, PA, Cat# Q350AR1A). After waiting for about 60 sec, the grid was plunge-frozen to liquid ethane using a ThermoFisher Mark IV vitrobot (ThermoFisher Scientific, Waltham, MA) with a blotting condition of 3.5 sec blot time, 0 blot force, and 100% humidity at 6°C. For E146Q-ATP and WT-AMP-PNP samples, 5 mM ATP or AMP-PNP was briefly incubated with 10 mM Mg^2+^ and ZnuBC for 60 sec at room temperature before blotting under the same conditions as the nucleotide-free samples.

Cryo-EM data for WT-AMP-PNP/Zn^2+^ state, E146Q-ATP/Zn^2+^ state, apo/Zn^2+^ state, and EDTA-treated/Zn^2+^-free state were collected at the LBMS facility at Brookhaven National Laboratory using a ThermoFisher Titan Krios 300 kV electron microscope equipped with a Gatan K3 camera and a BioQuantum energy filter. The physical pixel size used for all data collection is 0.666 Å. The beam size is 1.45 μm in diameter, covering the entire 0.6-μm hole. For each movie data frame, a total dose of 66 e^−^/Å^2^ was fractioned to 50 frames using the ThermoFisher data acquisition program EPU. All data sets were collected with an energy filter width of 15 eV, which was centered every 1 hr of data acquisition. Data collection statistics are listed in **Table S2**.

### CryoEM data processing

Fractioned movies were motion-corrected and averaged using MotionCorr2 ^38^ with a bin factor of 2. Averaged micrographs were further corrected by CTF estimation using Gctf ^39^. Micrographs with an estimated resolution better than 4.5 Å were selected for further use. Particles were initially picked in Relion3 ^40^ and cleaned up in cryoSPARC ^41^. Then the cleaned particles were used for training and particle picking using Kpicker ^42^.

For each data set, the picked particles were extracted at 256 pixels and binned to 64 pixels with a pixel size of 2.664 Å. We used cryoSPARC ^41^ for 2D class averaging to very generously select classes and particles 3D classification and refinement. With three classes each, selected particles were used for two cycles of *ab initio* reconstruction followed by 3D heterogeneous refinements with a particle size of 64 pixels (2.664 Å) in cryoSPARC. The particles belonging to classes with structural features were centered and re-extracted at 256 pixels and binned to 128 pixels with a pixel size of 1.332 Å. The re-extracted particles were used for 3D heterogeneous refinement using 3 or 4 classes. The process is repeated in cryoSPARC until there is no improvement in the refined map resolution as estimated by the gold-standard Fourier Shell Correlation (GS-FSC). There was no symmetry used during the iterative *ab initio* model generation and 3D heterogeneous refinements in cryoSPARC.

Particles from the best class or classes in cryoSPARC were auto-refined to convergence in Relion3 ^40^ followed by non-alignment 3D classification and local refinement without a symmetry (**Figure S3**). Particles from the best 3D class or classes with the best structural feature, as visualized in Chimera ^43^, were selected for non-uniform refinement and local refinement in cryoSPARC. With the WT-AMP-PNP/Zn^2+^, E146Q-ATP/Zn^2+^, and apo/Zn^2+^ states, C2 symmetry was justified to be appropriate based on GS-FSC (**Fig. S4**). All final particles are 128 pixels except for the ANP-PNP data which we extracted the final particles at 160 pixels. Local resolutions were estimated using BlocRes ^44^. Data processing and reconstruction statistics are listed in **Table S2**.

### Model building and refinement

To enhance the features for model building and visualization, the masked and filtered amps were processed by CryoFEM, which is a 3D convolutional neuron network (3D-CNN) model for local map sharpening and feature enhancement ^45^. The models were built in COOT ^46^ and refined in real space using PHENIX ^47^. The refined models were validated using Molprobity ^48^. The refinement statistics are listed in **Table S2**.

### Evaluation of transporter dynamics

All analyses for examining the dynamics of ZnuBC, including its potential transition to the inward-facing state, were performed using the *ProDy* API^26^. The Anisotropic Network Model (ANM) analysis ^49,50^ on ZnuBC was conducted by constructing a Hessian matrix (*3N × 3N*) for the tetrameric apo structure (ZnuBC) and the EDTA-treated structure (no ZSD) based on Cα coordinates. Explicit membrane ANM (ex MembrANM) ^27^, a method developed to incorporate lipid membrane in the ANM studies of membrane proteins embedded in an explicit membrane, was also performed on the apo structure to assess the effect of the membrane on protein dynamics. The membrane was constructed using the OPM server ^51^. The Hessian matrices were decomposed into ANM normal modes to extract the lowest frequency (softest) modes, which encapsulate global and cooperative motions. The first four ANM modes from the apo structure were reconstructed into 3D motions to visualize the individual mode motions (**Fig. S13**).

